# Practical considerations for accurate estimation of diffusion parameters from single-particle tracking in living cells

**DOI:** 10.1101/2025.06.12.659344

**Authors:** Aishani Ghosal, Yu-Huan Wang, Nguyen Nguyen, Laura Troyer, Sangjin Kim

**Author notes:** Corresponding author; (Electronic mail).

## Abstract

Advances in fluorescence microscopy have enabled high-resolution tracking of individual biomolecules in living cells. However, accurate estimation of diffusion parameters from single-particle trajectories remains challenging due to static and dynamic localization errors inherent in these measurements. While previous studies have characterized how such errors affect mean-squared displacement (MSD) analysis, practical guidelines for minimizing them during data acquisition and correcting them during analysis are still lacking. Here, we combine theoretical modeling and simulations to evaluate how exposure time and sampling rate influence the accuracy of MSD-based inference under fractional Brownian motion (FBM), a canonical model of anomalous diffusion. We demonstrate that decoupling exposure and sampling times enables escape from the error-prone regime, thus improving inference accuracy, and that incorporating an offset in nonlinear MSD fitting substantially improves the estimation of the anomalous diffusion exponent. We validate this framework using trajectories of cytoplasmic particles in *Escherichia coli*, recovering consistent diffusion parameters across multiple data sets. Our findings offer practical strategies to improve both experimental design and data analysis in single-particle tracking of live or synthetic systems.

## I. INTRODUCTION

Single-particle tracking (SPT) is widely used to investigate the local mechanical properties of living cells and biological fluids.^1,2^ These mechanical properties can be inferred by analyzing particle trajectories using various statistical measures.^3,4^ Among those, mean-squared displacement (MSD) is one of the primary analytical tools used to characterize diffusion. MSD quantifies the average distance traveled by a particle over a given time interval (lag time, *τ*) and is typically calculated by averaging squared displacements across multiple trajectories and time points as follows:

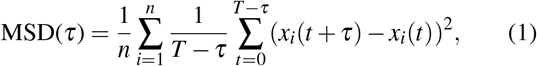

where *x*(*t*) denotes the particle’s position at time *t* during its evolution over the duration [0, *T*]. In this equation, the time average (TA) is performed over time points from 0 to *T* − *τ*, and the ensemble average (EA) is performed over *n* independent trajectories. The MSD increases with lag time (*τ*). The exact scaling relation is characterized with the diffusion coefficient (*D*) and the anomalous diffusion exponent (*α*) as follows:

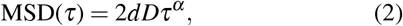

where *d* is dimensionality (e.g., *d* = 1 for one-dimensional diffusion). For normal diffusion (*α* = 1), the MSD increases linearly with lag time. In the case of subdiffusion, the MSD exhibits sublinear scaling with 0 < *α* < 1. Subdiffusive behavior has been reported across diverse biological systems, from *in vitro* networks^5–7^ to *in vivo* environments, such as bacterial cytoplasm and chromosomal loci.^8–12^ For example, Golding and Cox^9^ observed RNA-protein complexes exhibiting subdiffusive motion with *α* ≈ 0.7 in the cytoplasm of *E. coli* cells, and Weber et al.^10^ reported *α* ≈ 0.39 for the dynamics of chromosomal DNA loci in bacteria.

Characterizing anomalous diffusion is of great interest in soft matter and biophysics, as the value of *α* reflects interactions between the probe and its environment—such as crowding, binding, confinement, or viscoelasticity.^1,13^ Several models, including fractional Brownian motion (FBM) and continuous time random walk (CTRW), have been proposed to describe subdiffusive behavior.^14,15^ However, accurate interpretation of the underlying mechanisms remains challenging due to measurement artifacts intrinsic to SPT experiments. Localization errors—both static and dynamic—can distort the observed trajectories and bias the estimation of diffusion parameters from MSD.^16–19^ For example, static localization error alone can introduce apparent subdiffusive behavior in systems undergoing normal diffusion, highlighting the complexity of reliable MSD interpretation.^16,17^ Previous theoretical studies have explored how specific artifacts impact MSD.^16,19,20^ Notably, theoretical expressions for the expected MSD that account for both static and dynamic localization errors have been derived.^16,19^ Yet, a general framework for extracting true diffusion parameters from experimentally observed MSD remains lacking, particularly when imaging parameters, such as exposure time and frame rate, vary across experiments. In practice, exposure time (duration of light illumination) and frame interval (inverse of frame rate) are often selected empirically, without a clear understanding of their impact on MSD analysis. While prior work has examined the influence of illumination and camera integration,^21,22^ a systematic investigation of how these parameters jointly affect MSD analysis is still missing.

In this work, we address the challenge of recovering true diffusion parameters in the presence of static and dynamic localization artifacts in SPT data. We follow the theoretical formalism derived by Backlund et al^19^ for FBM and examine how inferred *D* and *α* are influenced by static and dynamic localization errors under various imaging conditions. Finally, we apply these theoretical insights to extract diffusion parameters from published and newly acquired SPT data. This work provides an essential tool for the biophysics community, improving the reliability of MSD analysis in probing complex cellular environments.

The paper is structured as follows: Section II presents the theoretical background and simulation methods. Section III describes the SPT imaging protocols and their limitations, and proposes refined imaging conditions. Section IV outlines improved approaches for extracting accurate diffusion parameters, validated using using simulated data. Section V presents the application of these methods to experimental data, and Section VI concludes the study.

## II. MODEL

Among several theoretical models for subdiffusion, FBM has been shown to describe subdiffusive dynamics of biomolecules in cells in multiple cases.^23^ In addition, Backlund et al. have derived an MSD equation incorporating both static and dynamic localization errors.^19^ Therefore, we focus our analysis on FBM and extend previously derived analytical expressions to various imaging protocols.

In essence, the MSD of FBM originates from the two-point position correlation function (at time *t*_1_ and *t*_2_), which follows a power law with respect to the time difference:^24^

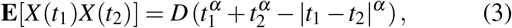

where **E** denotes ensemble average. In actual SPT experiments, particle positions are recorded at discrete time points. Therefore, we hereafter denote time in terms of frame indices. The frame index *k* can be converted to time by multiplying the frame interval, *γ* (i.e., the inverse of the frame rate or sampling rate). We note that this frame interval can differ from the light exposure time (*t*_*E*_) depending on the imaging protocol. In the case of continuous exposure, the frame interval (*γ*) is equal to the light exposure time (*t*_*E*_), as shown in Fig. 1(a). In the case of time-lapse imaging mode, *γ* is larger than *t*_*E*_ (Fig. 1(a)); the camera shutter is closed between frames. Additionally, *t*_*E*_ refers to the light exposure time, which may differ from the camera shutter open time in the case of stroboscopic imaging.^25^ In our discussion, the light exposure time (not the camera shutter open time) is relevant because it defines the period during which photons are emitted. We model both continuous exposure and time-lapse protocols, schematically shown in Fig. 1(a), to study their effects on MSD and identify suitable imaging parameter regimes that result in lower MSD error, as illustrated in Fig. 2(c,f) and Fig. 3(e-h).

**FIG. 1.**
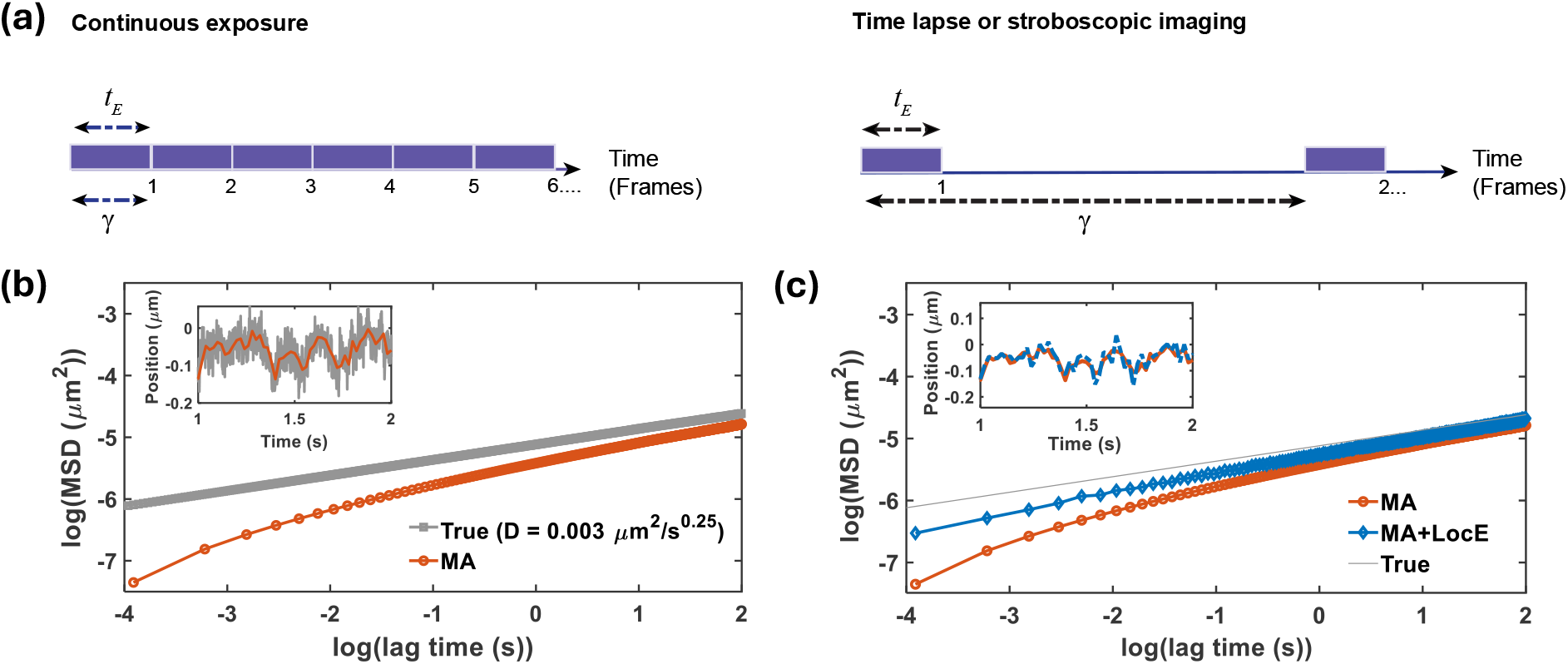
Schematic presentation of imaging protocols and simulated trajectories of a particle exhibiting FBM. (a) Experimental protocols based on continuous exposure (left) and time-lapse imaging (right). Fluorescence signal is acquired during *t*_*E*_, and images are taken at the interval of *γ*. In continuous exposure, *t*_*E*_ = *γ*, while in time-lapse mode, *t*_*E*_ < *γ*. (b-c) Effects of dynamic (MA) and static localization (LocE) errors on MSD. In (b), grey and red markers represent the true and MA-applied EA-TA MSDs, respectively. (Inset) Grey line shows the simulated 1D trajectory of FBM sampled every 1 ms. Red line shows the trajectory with MA applied (*t*_*E*_ = 20 ms). In (c), red and blue markers represent EA-TA MSDs with MA only and with both MA and LocE, respectively. The grey line corresponding to the true EA-TA MSD (same as in (b)) is shown for reference. (Inset) Red and blue lines show 1D FBM trajectories with MA only and with both MA and LocE, respectively. The parameters used for the simulation are as follows: *D* = 0.003*µm*^2^/s^0.25^, *α* = 0.25, *t*_*E*_ = 20 ms, and *σ* = 20 nm. log refers to the natural logarithm.

**FIG. 2.**
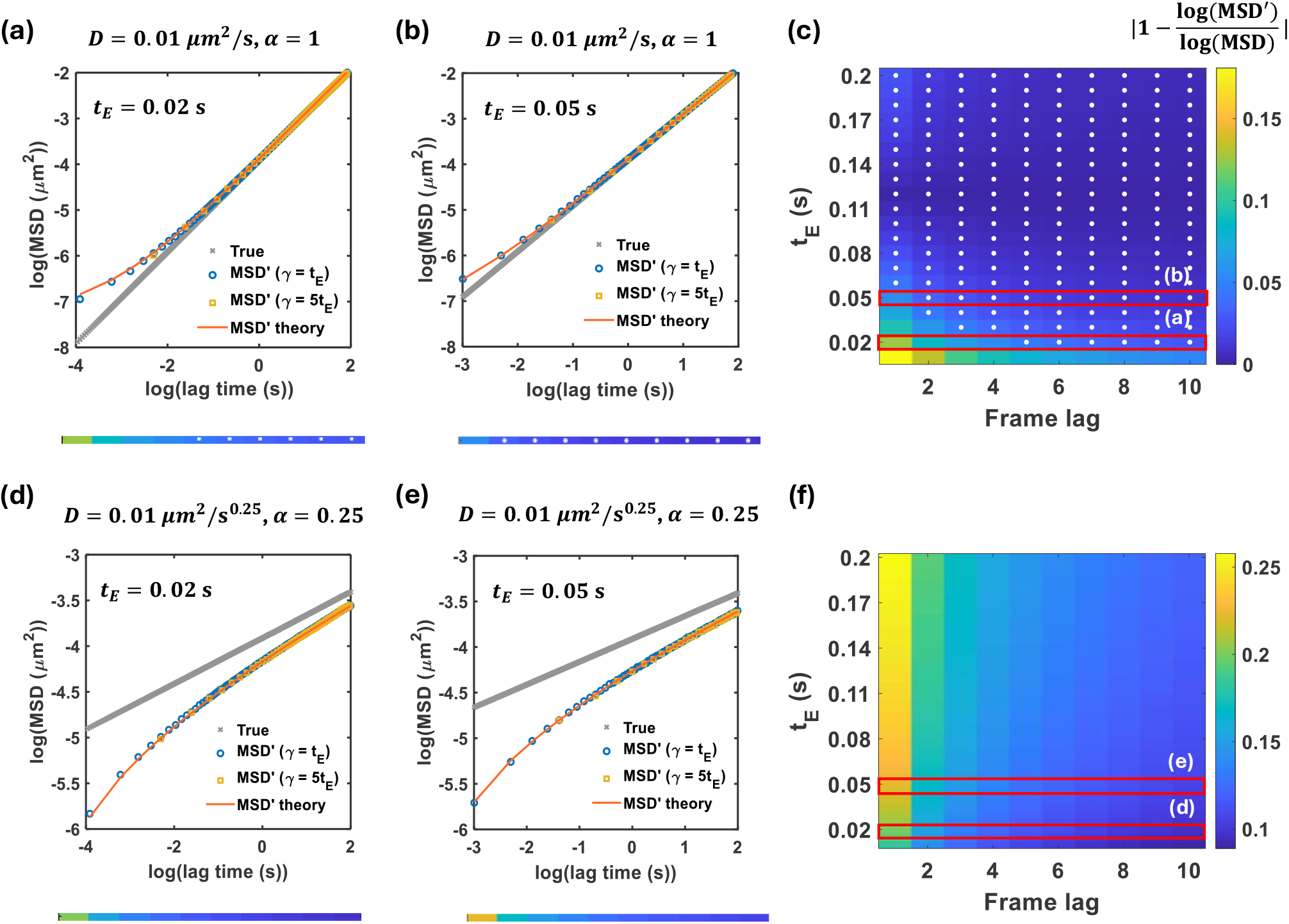
Comparison between observed MSD and true MSD across the parameter space of *t*_*E*_ and *γ*. The observed MSD (MSD*′*) includes both MA and LocE, and the error relative to the true MSD (MSD) is calculated as |1− log(MSD*′*)*/*log(MSD) |. (a,b,d,e) Example EA-TA MSD curves in the absence and presence of MA and LocE, under continuous exposure and time-lapse imaging protocols. The yellow solid line corresponds to the theoretical prediction from Eq. (8). The remaining simulation parameters are *D* = 0.01*µm*^2^/s^0.25^, *σ* = 20 nm; *α* = 1 (a-b) and 0.25 (d-e); *t*_*E*_ = 0.020 s (a,d) and 0.050 s (b,e). Color bars below each plot display the relative error corresponding to each set of parameters. (c, f) Colormaps of relative error in the presence and absence of MA and LocE, plotted as a function of *t*_*E*_ and frame lag. The white dots denote the relative error ≤ 5% for the indicated parameter values (*α* = 1 in (c) and 0.25 in (f)).

**FIG. 3.**
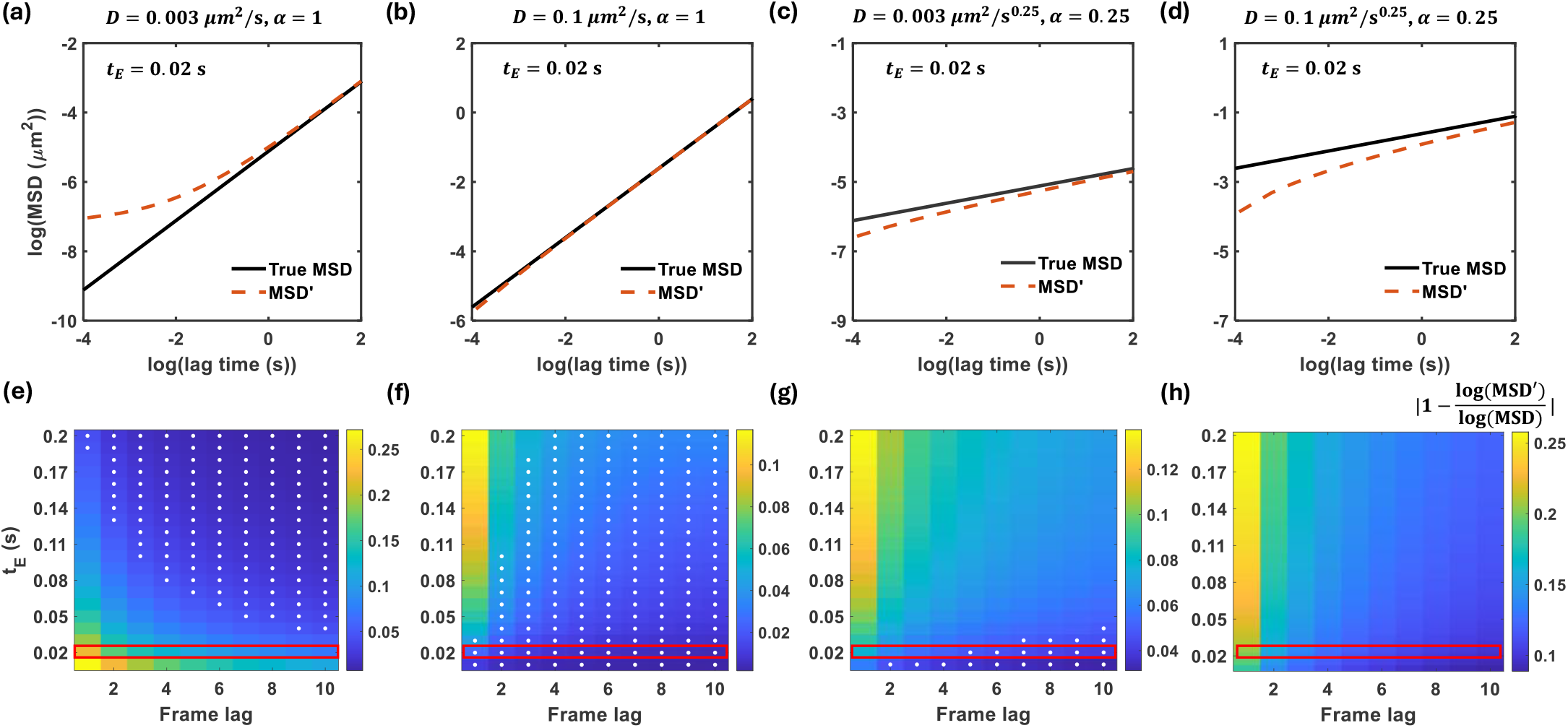
Effect of underlying diffusion dynamics on the relative error in the observed MSD. Upper panels show the MSD in the presence (red dotted line) and absence (black solid line) of the microscopy effects for diffusion parameter values (a) *D* = 0.003*µm*^2^/s, *α* = 1; (b) *D* = 0.1*µm*^2^/s, *α* = 1; (c) *D* = 0.003*µm*^2^/s^0.25^, *α* = 0.25; (d) *D* = 0.1*µm*^2^/s^0.25^, *α* = 0.25. The lower panel shows the relative error between the MSDs in the presence and absence of the microscopy artifacts, and the white dots indicate the relative error less than or equal to 5%. Red boxes highlight the relative errors corresponding to the upper panel (*t*_*E*_ = 20 ms).

### A. Theory

EA-TA MSD defined in Eq. (1) as a function of lag time is expressed as a function of frame lag *n* (or the number of frames spanning the lag),

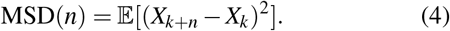

Here, 𝔼 stands for ensemble averaging and time averaging over *k. X*_*k*_ represents the true particle position at the *k*th frame, measured in the absence of measurement artifacts. Lastly, MSD(n) denotes the true MSD (without measurement artifacts).

This EA-TA MSD is characterized with *D* and *α* as follows:

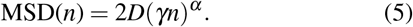

This expression corresponds to MSD in one dimension (1D), and the 3D case is obtained by summing the MSDs of three independent 1D processes (Eq. (2)).

During light exposure of *t*_*E*_, particles continue to move, and only the averaged position is recorded in each frame. This averaging is referred to as motion averaging (MA), also called dynamic localization error in the literature, because this effect arises from particle dynamics.^16,26^ In addition, due to photon statistics, the position of a particle is measured with finite localization precision.^27^ This contributes to the static localization error (LocE). Namely, during *t*_*E*_ for the *k*th frame, the particle emits *p* number of photons, where *p* follows a Poisson distribution with mean 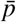. Expressing the particle position at the time of photon *i* emission as 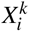, we add a random variable 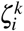 to account for static localization error. This random variable is modeled as Gaussian-distributed random noise with a standard deviation, *s*_0_, which is determined by the width of the microscope’s point spread function. The estimated (measured) position of the particle at the *k*th frame is then given by:

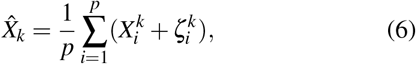

where the hat denotes the estimated quantity. This expression includes both static and dynamic localization errors.

Next, the right-hand side of Eq. (4) is rewritten using the observed trajectories as follows:

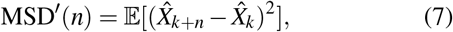

where MSD^*′*^(n) denotes the observed MSD including measurement artifacts. We followed the detailed derivation by Backlund et al^19^ for 1D MSD, and we extended it to accommodate time-lapse and stroboscopic imaging protocols (*γ* ≥ *t*_*E*_, as shown in Fig. 1(a)). The final expression is given by:

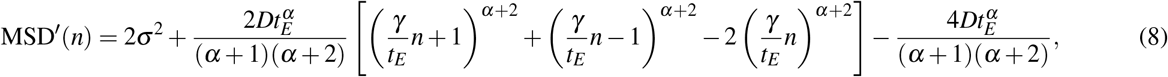

where *n* = 1, 2, 3, *· · ·*.

According to the derivation by Backlund et al., the first term of Eq. (8) is not simply the localization error of an immobile particle (*σ*_0_), yet it also includes additional spreading of photons due to particle motion—an image blurring effect.^19,20,26^ We note that this effect is distinguished from motion averaging effect (MA) during *t*_*E*_. *σ* ^2^ is related to 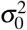, the localization error of a stationary particle as follows:^19^

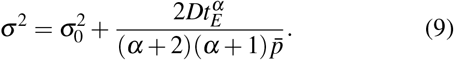

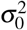 equals 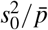, and 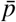 would increase with larger *t*_*E*_. Additionally, if many photons are collected during a frame (as 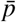 approaches infinity), *σ* approaches *σ*_0_.

The primary motion averaging effect is explained by the last term of Eq. (8). The middle term (within the square brackets) also contains contributions from motion averaging. By expressing the middle term as an infinite series and considering only the first few non-zero terms, it can be shown that the leading term still scales with *n*^*α*^. The non-leading terms are most significant when *n* = 1.^19^ This feature is later used in Sec. IV for the nonlinear fitting of MSD.

In the case of normal diffusion (*α* = 1), Eq. (8) simplifies as previously shown:^16,20,21^

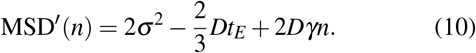

We show in Sec. II B that the intrinsic measurement artifacts can contribute a net positive or negative offset to the true MSD, depending on the magnitude of the diffusion parameters and imaging conditions.

### B. Simulation

Trajectories of FBM dynamics were simulated based on the power-law time-dependent position correlation given in Eq. (3) for a set of diffusion parameters. The standard deviation of the position covariant matrix was considered as the step size and used to update the particle’s position:

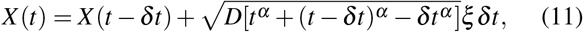

where *δt* is the sampling time of the simulation, *ξ* is a random noise term with zero mean and delta-correlated variance, and the initial condition is *X* (0) = 0. *X* (*t*) and *X* (*t* − *δt*) represent the true particle positions at time *t* and an earlier time *t*− *δt*, respectively (See Appendix A for the details of the simulation).

After generating a 10 s-long time-series of *X* (*t*) with a sampling time *δt* of 1 ms, the positions were averaged over a time window of *t*_*E*_ to account for the motion averaging effect (Fig. 1(b), inset). Static localization error was then added to each time-averaged position at frame *k* as follows:

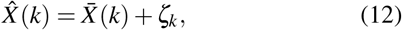

where *ζ*_*k*_ is a random variable drawn from a Gaussian distribution with variance 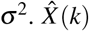 denotes the observed trajectory, which includes both the motion averaging effect 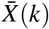 and the static localization error (Fig. 1(c), inset).

As shown in Fig. 1(b-c), microscopy effects influence not only the measured trajectories 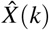 but also affect the MSD accordingly. When the motion averaging effect is added, the log-log MSD curve bends downward (Fig. 1(b)). In contrast, the static localization error has the opposite effect, producing an upward shift (Fig. 1(c)). These opposing effects can also be anticipated from Eq. (8).

## III. EFFECT OF IMAGING CONDITIONS ON THE OBSERVED MSD

As static and dynamic localization errors have antagonizing effects, one can determine the optimal imaging conditions by setting the terms related to these effects equal in Eq. (8). However, the solution depends on the unknown ground-truth values of diffusion parameters. This section discusses how to tune imaging conditions to reduce the microscopy artifacts that propagate into the observed MSD.

As we consider constant *σ* = 20 nm, varying the imaging conditions simply refers to adjusting *t*_*E*_ and *γ* (See Appendix C for a relaxation of the assumption on *σ*). Given fixed diffusion parameters (*D* and *α*), decreasing *t*_*E*_ and increasing *γ* reduce the motion averaging effect due to changes in the correlation between frames.

To demonstrate the impact of imaging conditions (*t*_*E*_ and *γ*), we initially used two examples: *D* = 0.01*µm*^2^/s for normal diffusion and *D* = 0.01*µm*^2^/s^0.25^, *α* = 0.25 for subdiffusion (Fig. 2). The simulated MSD incorporating MA and LocE (or the observed MSD denoted as MSD^*′*^) showed good agreement with the theoretical prediction from Eq. (8), validating our simulation approach. We tested how continuous exposure and time-lapse imaging protocols affect MSD^*′*^ for a given *t*_*E*_ (20 ms in Fig. 2(a,d) and 50 ms in Fig. 2(b,e)). For a given *t*_*E*_, the two imaging protocols produced the same degree of MA in the MSD^*′*^, but at different time lags.

In the case of normal diffusion *D* = 0.01*µm*^2^/s, the MA effect was smaller than the LocE effect when the shortest *t*_*E*_ = 20 ms was used, resulting in an upward-bending log-log MSD curve at short lag times (Fig. 2(a)). The deviation of MSD^*′*^ from the true MSD (which does not have either MA and LocE effect) was evident only at the short time lags - for example, below log(lag time) ≈−2 in Fig. 2(a). When a longer *γ* was used in time-lapse mode (e.g., *γ* = 100 ms instead of 20 ms), MSD^*′*^ was closer to the true MSD because it started from a regime where MSD^*′*^ had already approached the true MSD (Fig. 2(a)).

When *t*_*E*_ = 50 ms, the MA effect was larger than the case of *t*_*E*_ = 20 ms, such that the difference between MSD^*′*^ and true MSD was smaller (Fig. 2(b)). For other values of *t*_*E*_, we calculated the difference between the observed MSD and the true MSD using the relative error between log(MSD^*′*^) and log(MSD), as the MSD is analyzed in log-log space (Fig. 2(c)). Specifically, we calculated |1-log(MSD^*′*^) /log(MSD)|. The colormap shows how the relative error varies as a function of *t*_*E*_ and frame lag. Each row represents the relative error calculated from log(MSD^*′*^) at different frame lags for continuous exposure with the specified *t*_*E*_. The white dots indicate the relative error less than or equal to 5%. Fig. 2(c) shows that, for a given *D* value for normal diffusion, the difference between MSD^*′*^ and MSD is small across all frame lags when *t*_*E*_ is greater than 60 ms (white dots). We note that while we assumed a constant *σ* = 20 nm, increasing *t*_*E*_ can raise 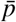 and the image blurring term, *Dt*_*E*_, in Eq. (9). They have opposing effects on *σ* according to Eq. (9). In fact, when *t*_*E*_ becomes very large, its effect on *σ* becomes non-negligible, and *σ* ≈ 20 nm does not hold anymore. We will discuss *σ* as a function of *t*_*E*_ in Appendix C.

In the case of subdiffusion (*D* = 0.01*µm*^2^/s^0.25^), the MA effect was larger than the LocE effect even at *t*_*E*_ = 20 ms, causing the MSD^*′*^ to bend downward at early lag times (Fig. 2(d)). Interestingly, MSD^*′*^ did not approach the true MSD even at later time lags. The relative error between log(MSD^*′*^) and log(MSD) remained over 5% (i.e. white dots do not appear in the color bar). For longer *t*_*E*_ = 50 ms, the MA effect increased, and the error became even larger (Fig. 2(e-f)). These examples illustrate that the upward or downward curvatures of the observed MSD in log-log space is influenced by the combined effect of MA and LocE, where the former is strongly dependent on the ground-truth *D* and *α*, as well as on the chosen imaging conditions (*t*_*E*_ and *γ*).

Using the theoretical prediction of MA and LocE effects on MSD^*′*^ (Eq. (8)), we explored the impact of *t*_*E*_ across different regions of the diffusion parameter space defined by *D* and *α* (Fig. 3). When the lowest *t*_*E*_ of 20 ms was used, MSD^*′*^ exhibited either upward or downward curvature depending on the underlying diffusion dynamics (Fig. 3(a-d)). Notably, when MSD^*′*^ was dominated by LocE (Fig. 3(a)), increasing *t*_*E*_ helped bring MSD^*′*^ close to the true MSD at early time lags (Fig. 3(e)). In contrast, when MSD^*′*^ was close to the true MSD due to a coincidental cancellation of MA and LocE effects (Fig. 3(b-c)), increasing *t*_*E*_ amplified the MA contribution and thus increased the relative error (Fig. 3(f-g)). Finally, when MSD^*′*^ was already dominated by MA at the lowest *t*_*E*_ (Fig. 3(d)), further increasing *t*_*E*_ did not improve agreement with the true MSD (Fig. 3(h)).

This analysis helps predict the optimal *t*_*E*_ and *γ* for imaging, even without knowing the ground-truth *D* and *α*. If the MSD^*′*^ measured at the lowest *t*_*E*_ shows an upward curvature in log-log space (Fig. 3(a)), a larger *t*_*E*_ may improve accuracy. Conversely, if MSD^*′*^ curves downward, as in Fig. 3(d), adjusting *t*_*E*_ does not help. Additionally, even when MSD^*′*^ is curved at early time lags (in a log-log scale), it often follows a straight line at later time lags, suggesting that using MSD^*′*^ at longer time lags can mitigate errors. Experimentally, if trajectories are sufficiently long, one may exclude short time lags and use only long-lag MSD values. However, this is not always feasible, particularly when trajectories are short due to photobleaching. In such cases, rather than collecting many unnecessary frames at short time intervals, it is more practical to employ time-lapse acquisition with a large *γ*. When a large *γ* is used, the initial MSD^*′*^ (i.e., the shortest time lag) may already fall within the ≤ 5% relative error regime. For example, in Fig. 3(c,g), using *γ* = 5*t*_*E*_ results in all MSD^*′*^ points remaining within the ≤ 5% error range.

We note that in other examples (as in Fig. 3(d)), it may not be possible to achieve MSD^*′*^ ≤ 5% simply by adjusting *t*_*E*_ and *γ*. Furthermore, using a large frame interval in time-lapse imaging may not be always practical. A large *γ* can lead to incorrect particle linking between frames and introduce confinement effects due to cell boundaries. In the next section, we introduce a MSD fitting method in which MSD^*′*^ is used *as is* to extract diffusion parameters.

## IV. DIFFUSION PARAMETER EXTRACTION FROM SIMULATED DATA

In this section, we investigate methods to extract ground-truth diffusion parameters from the observed MSD (MSD^*′*^), based on simulated FBM trajectories as detailed in Sec.II B. Prior studies proposed estimation based on maximum likelihood methods^28^ and linear fitting of MSD in log-log space.^17^ However, Eq. (8) suggests that static and dynamic localization errors introduce an offset to the ground-truth MSD. Therefore, conventional linear fitting may not yield accurate parameter estimates. We compared fitting EA-TA MSD using either a linear or a nonlinear model to extract the diffusion parameters. In both cases, we used initial MSD points for the fitting in log-log space. Although these points from short time lags are more affected by microscopy artifacts, as shown in Fig. 2 and Fig. 3, they are statistically more robust and are commonly used in experimental MSD analysis.^29^

### A. Linear fitting

EA-TA MSD vs. lag times were plotted on a log-log scale, and a linear fit was performed following Eq. (5) as follows:

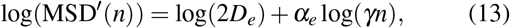

where the slope is *α*_*e*_ and the y-intercept is log(2*D*_*e*_). *D*_*e*_ and *α*_*e*_ denote the inferred diffusion coefficient and diffusion exponent, respectively.

By fitting MSD^*′*^ with Eq. (13) for two different imaging protocols, we found that the extracted parameters (*D*_*e*_ and *α*_*e*_) deviate from the ground-truth values (*D* and *α*) (Fig. 4(a-b)). This deviation arises from the omission of the constant offset term, which accounts for measurement artifacts. However, we found that *α*_*e*_ obtained from time-lapse imaging (*γ* = 5*t*_*E*_) is closer to the ground truth than that from continuous exposure (*γ* = *t*_*E*_), due to its ability to access larger lag times with the same number of frames. As shown in Fig. 2 and Fig. 3, larger lag times reduce the difference between MSD^*′*^ and the true MSD.

**FIG. 4.**
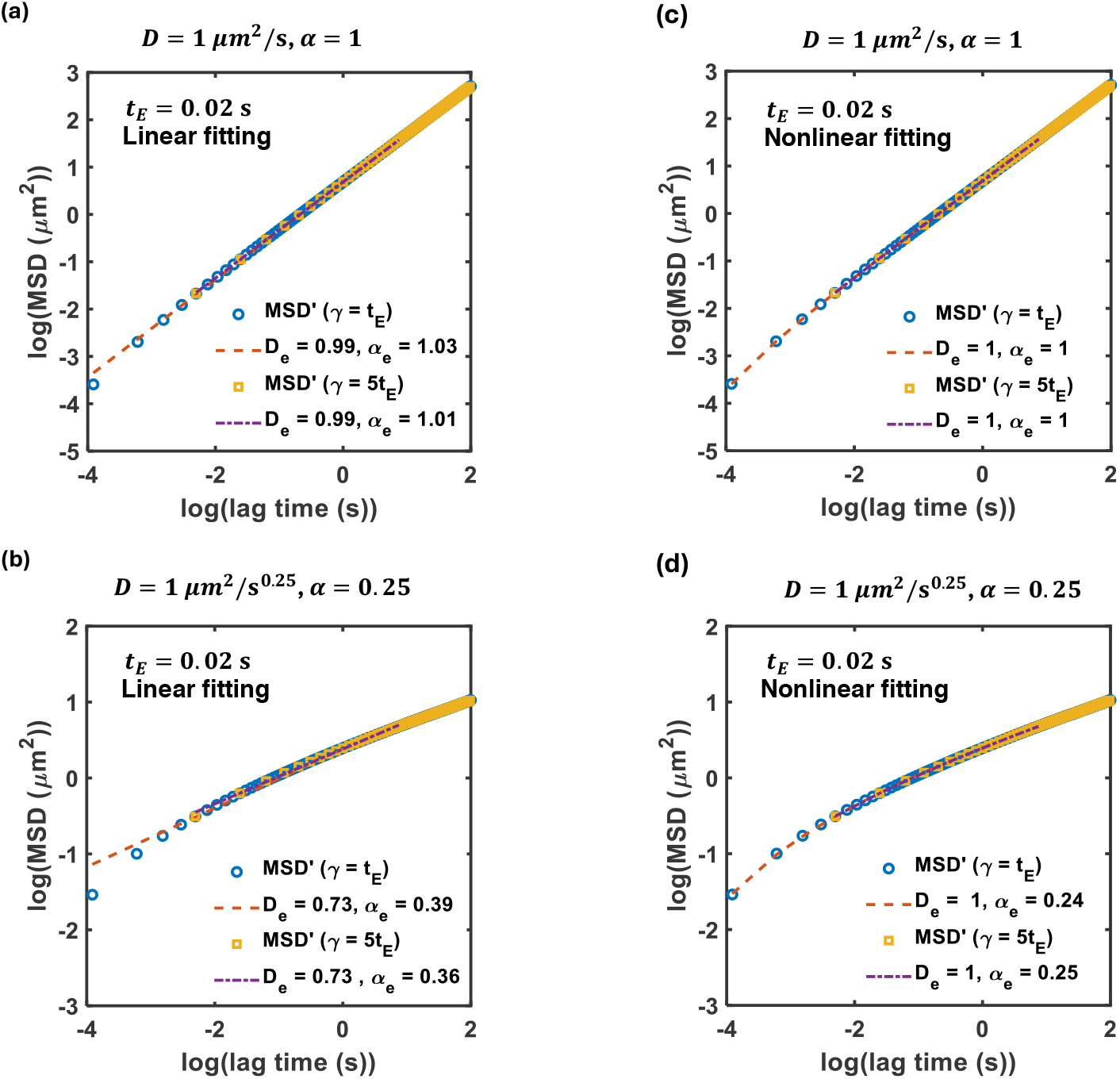
Extraction of diffusion parameters by fitting simulated EA-TA MSD and lag times in log-log space. (a, b) with linear model given by Eq. (13) and (c, d) with nonlinear model given by Eq. (14). Legend shows the inferred diffusion parameters (*D*_*e*_ and *α*_*e*_) for the following ground truth parameter values: (a) *D* = 1 *µm*^2^/s, *α* = 1; (b) *D* = 1 *µm*^2^/s^0.25^, *α* = 0.25; (c) *D* = 1 *µm*^2^/s, *α* = 1; (d) *D* = 1 *µm*^2^/s^0.25^, *α* = 0.25; together with *t*_*E*_ = 20 ms and *σ* = 20 nm.

### B. Nonlinear fitting

EA-TA MSD vs. lag times were plotted on a log-log scale, and a nonlinear fit was performed as follows:

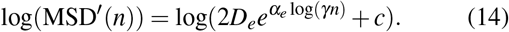

The offset term *c* in the equation is analogous to the terms related to the static and dynamic localization errors in Eq. (8). As shown in Fig. 4(c-d), both continuous and time-lapse imaging protocols perform well in estimating the true diffusion parameters (*D* and *α*). We note that the nonlinear fitting model Eq. (14) omits several high order terms in Eq. (8) to allow the fitting feasible, yet the inferred diffusion parameters remain close to the ground-truth values. In the next section, we demonstrate that this nonlinear fitting approach is also applicable to experimental data.

## V. DIFFUSION PARAMETER EXTRACTION FROM EXPERIMENTAL DATA

We applied the nonlinear fitting method to extract diffusion parameters from experimental data. As test cases, we used previously published MSD data sets from tracking RNA-protein complexes^12^ and chromosomal loci^30^ in *E. coli*.

The RNA-protein complex consisted of MS2-GFP fusion proteins bound to mRNA containing 96 tandem repeats of MS2 binding sequence.^9^ In the study of Lampo et al,^12^ SPT was performed at a frame interval *γ* of 1 s, and trajectories were analyzed in 1D along the long axis of the cell. They reported ergodic subdiffusive behavior with *α* ≈ 0.54.^12^ When we replotted their MSD data, we observed a subtle upward curvature at the early time lags, which, as discussed earlier, can result from microscopy artifacts (Fig. 5(a)). Conventional linear fitting yielded *D*_*e*_ = 0.0018 *µm*^2^/s^0.56^, *α*_*e*_ = 0.56, consistent with the original report. However, nonlinear fitting produced *α* of 0.67. Interestingly, this *α* value is close to that obtained from linear fitting of a data set acquired with a longer sampling interval (*γ* =1 min) in previous studies.^9,12^ This result is consistent with our finding that large *γ* can improve recovery of true diffusion parameters.

**FIG. 5.**
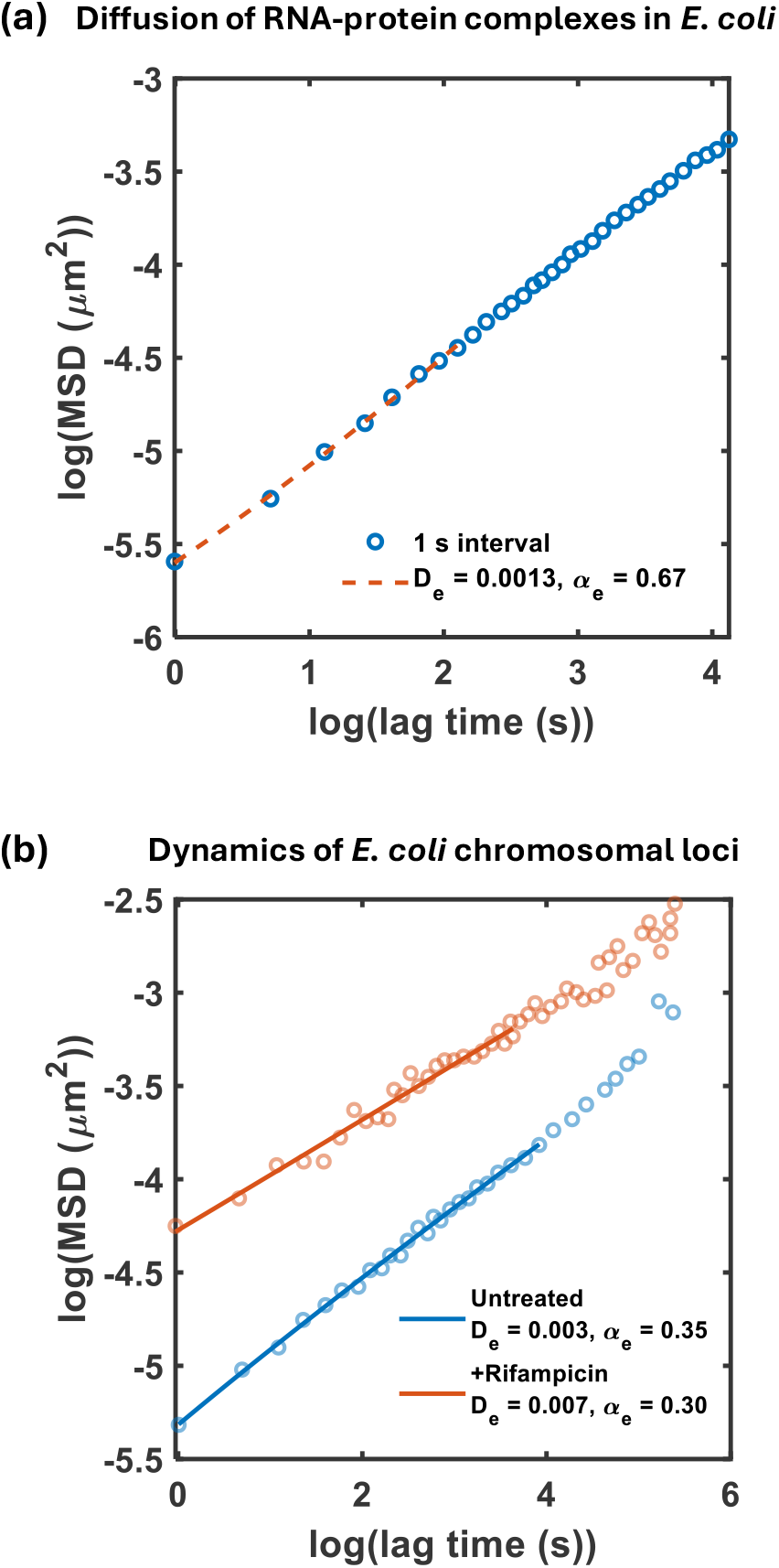
Extraction of diffusion parameters from experimental EA MSD using the nonlinear model. (a) Log-log MSD vs lag times for the motion of cytoplasmic RNA-protein complexes in *E. coli*.^12^ (b) Log-log MSD vs lag times for the motion of chromosomal loci in *E. coli*, with or without antibiotic treatment.^30^ Data points were extracted from published figures using Webplotdigitizer.^31^ Symbols are quantified from displacements along the long axis of the cell (1D), and lines correspond to nonlinear fits.

Chromosomal loci dynamics are well characterized by FBM.^10,19^ We captured the EA MSD of the chromosome loci reported in an earlier study.^30^ In the study, loci were visualized using GFP-tagged ParB proteins bound to a *parS* site on the chromosome, and displacements along the cell’s long axis were used for 1D MSD analysis. Nonlinear fitting of the MSD yielded *D*_*e*_ = 0.003 *µm*^2^/s^0.35^, *α*_*e*_ = 0.35, comparable to *α*_*e*_ = 0.38 from linear fitting.

In the same report,^30^ *E. coli* cells were treated with 100 *µ*g/mL rifampicin, an antibiotic that inhibits transcription^32^, for 30 min to deplete cellular RNA.^33^ Under this condition, the chromosomal loci exhibited larger displacements compared to those in untreated cells (Fig. 5(b)). Nonlinear fitting of the MSD yielded *D*_*e*_ = 0.007 *µm*^2^/s^0.30^, *α*_*e*_ = 0.30. Linear fitting also gave *α*_*e*_ = 0.30. The agreement between the two fitting methods reflects the fact that the MSD curve does not exhibit significant curvature in log-log space (Fig. 5(b)).

Next, we applied the fitting methods to our own experimental data obtained by tracking endogenous proteins in *E. coli* at a low *t*_*E*_ = *γ* of 20 ms (Fig. 6).^34^ The proteins were fused to a photo-convertible fluorescent protein mEos3.2.^35^ EA-TA MSD analysis of the SPT data revealed slightly lower *α* values by the nonlinear fitting method (*α* ≈ 0.61 − 0.76) compared to the linear fitting method (*α* ≈ 0.71 − 0.76). The small difference was found when the observed MSD is bent slightly downward (e.g., cytoplasmic RNase E (1-529) and LacZ).

**FIG. 6.**
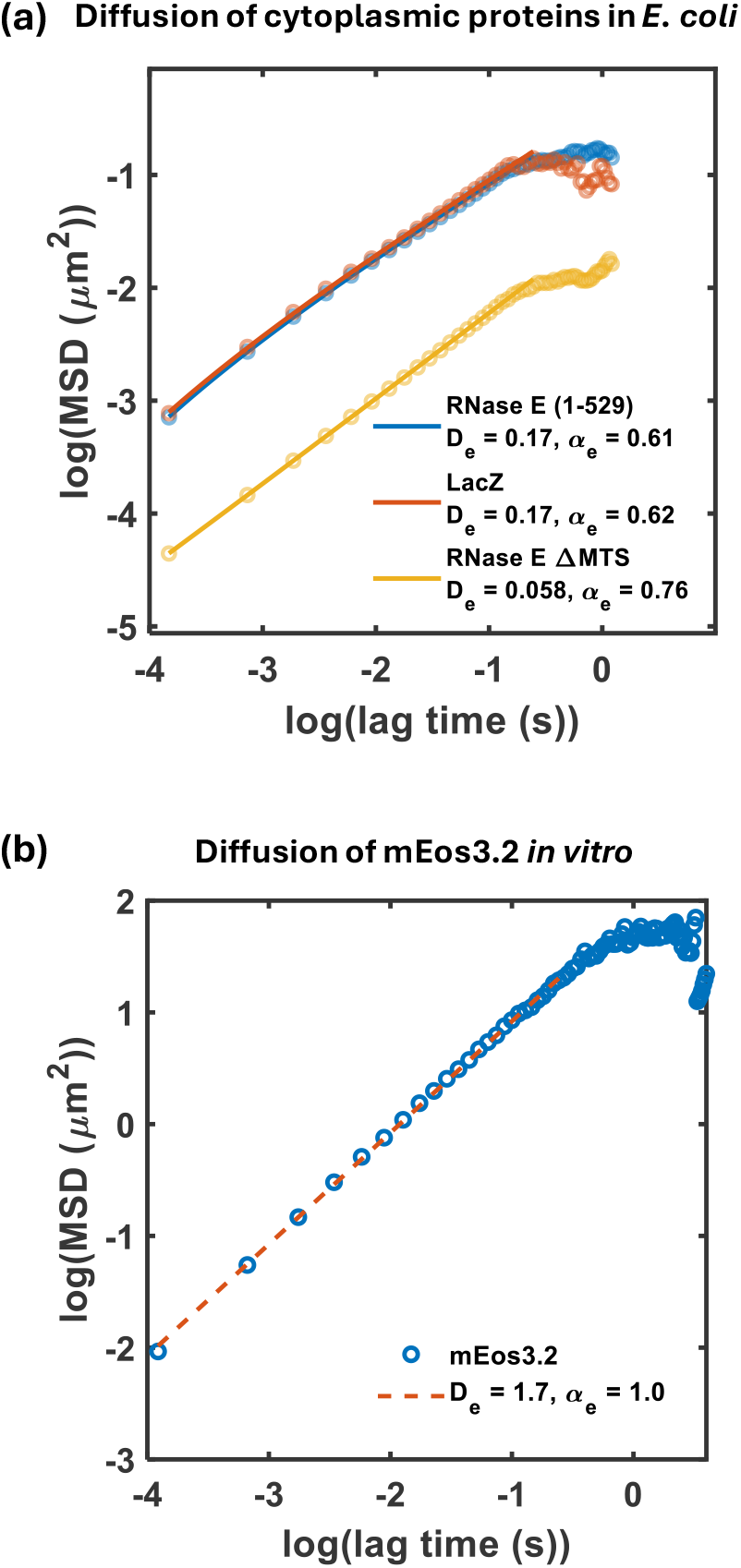
Extraction of diffusion parameters from experimental 2D EA-TA MSD acquired from continuous exposure of *t*_*E*_ = 20 ms. (a) Experimentally acquired MSD of mEos3.2-fused cytoplasmic proteins (LacZ and two types of cytoplasmic RNase E mutants) in *E. coli*. (b) Experimentally acquired MSD of mEos3.2 diffusion *in vitro*.

To test whether *α* < 1 is an intrinsic *in vivo* property, we purified the same fluorophore (mEos3.2) and tracked its motion *in vitro* under identical imaging conditions (*t*_*E*_ = *γ* = 20 ms). Fitting its MSD revealed *α* ≈ 1 *in vitro* using both linear and nonlinear methods (Fig. 6(b)). This indicates that the subdiffusive behavior observed *in vivo* arises from intracellular constraints. We also note that the *α* ≈ 0.6 − 0.7 values obtained from nonlinear fitting are consistent across distinct cytoplasmic probes (RNA-protein complexes shown in Fig. 5(a) and endogenous proteins shown in Fig. 6(a)), despite differences in *D* and imaging conditions. This convergence supports the idea that subdiffusion is a robust feature of the cytoplasmic environment in *E. coli*.^9^

## VI. CONCLUSION

SPT experiments have inherent measurement errors due to motion averaging and finite photon statistics in each frame. In this study, we examined how these intrinsic errors influence diffusion parameter estimation from SPT data. Using theoretical models, simulations, and experimental data analysis, we showed that these errors introduce systematic deviations in the observed MSD, particularly at short lag times. Motion averaging and static localization error have opposing effects on MSD curvature in log-log space, and their relative impact depends on the underlying diffusion parameters (*D* and *α*) and imaging conditions—specifically exposure time *t*_*E*_ and frame interval *γ*.

We identified parameter regimes that minimize MSD distortion and proposed diagnostic guidelines based on MSD curvature and lag-time dependence. Notably, larger *γ* and appropriate *t*_*E*_ values reduce the influence of measurement artifacts. We further assessed the performance of linear versus non-linear fitting methods for extracting *D* and *α*. While linear fitting failed to account for offset terms arising from measurement artifacts, nonlinear fitting, based on a simplified form of the theoretical MSD expression, yielded more accurate parameter estimates. The application to both published and newly acquired experimental data demonstrated the utility of this method. Specifically, we confirmed that *E. coli* cytoplasmic proteins exhibit subdiffusive behavior *in vivo*, whereas the same fluorophore displays normal diffusion *in vitro* under identical imaging conditions, reinforcing the biological significance of anomalous diffusion.

These results establish practical guidelines for minimizing error in SPT-based diffusion measurements and improve the reliability of parameter inference under complex experimental conditions.

## ACKNOWLEDGMENTS

We thank Maggie Liu for their contributions in the early phase of this work and the members of Kim lab for critical reading of the manuscript. This work was supported by the NSF Center for Physics of Living Cells (1430124), NSF Science and Technology Center for Quantitative Cell Biology (2243257), and NIH (R35GM143203).

## DATA AVAILABILITY STATEMENT

The datasets generated or analyzed during the current study are included in this manuscript.

## APPENDIX A: SIMULATION METHOD

To generate the step sizes for the simulated trajectory, we began by decomposing the position covariance matrix, the elements of which are given in Eq. (3). We used the Cholesky decomposition of this covariance matrix via the `MATLAB`^36^ function chol. Since the matrix is symmetric, only the lower triangular part was decomposed. The result was then multiplied by a normally distributed random vector to generate FBM trajectories. Next, we added motion averaging and static localization error and computed EA-TA MSD. The simulation used a sampling time of 1 ms, with a total of 10,000 steps, corresponding to 10 s. After time averaging, the MSD values were further averaged over 10,000 independent trajectory realizations. The resulting MSD^*′*^ values, computed for different diffusion parameters, are shown in Fig. 1(b-c) and Fig. 2.

## APPENDIX B: EXPERIMENTAL METHODS

Linear and nonlinear fitting was performed using `MATLAB` function fit in log-log space. Fitting in log space weighs early time points more heavily and enhances parameter sensitivity in that regime. Cytoplasmic protein tracking was performed in our previous study and details of the experiments can be found in the paper^34^. The *in vitro* data has not been published elsewhere. The expression and purification of the protein was described previously^34^. We diluted the protein to a final concentration of 36 pM in 94.95% glycerol. The protein mixture was dropped on a cleaned glass slide with an imaging spacer (Grace Bio-Labs, GBL654002) and sealed with a clean glass coverslip. To help find the imaging focal plane, we added Cy5-labeled DiagNano Fluorophore Labeled Gold Nanoparticles (CD Bioparticles, GFL-5) at 3.3 *µ*M concentration as well.

We took fluorescence images using a Ti-2 Nikon microscope equipped with a TIRF objective 100x/1.49 NA (Nikon), a 4-color laser launch (iChrome MLE-LFA-NI1, Toptica Photonics), and an EM CCD camera (Andor iXon Ultra). In this microscope setup, *γ* = 20 ms is the lower limit of the camera frame interval when the full chip is used. A 640-nm laser was used to image Cy5-labeled nanoparticles. After finding the focal plane, we used a 405-nm laser of 2.24 W/cm^2^ to photoactivate mEos3.2 and switched to a 561-nm laser to excite the photo-converted mEos3.2 molecules at an intensity of 3.06 W/cm^2^. Images were acquired every 20 ms via continuous exposure, the same way we performed SPT of proteins in *E. coli*.^34^

## APPENDIX C: STATIC LOCALIZATION ERROR OF MOVING PARTICLES

The first term in Eq. (8), *σ* ^2^, represents the static localization error arising from finite photon statistics. It reflects the localization precision based on the diffraction-limited image of fluorescent particles. In our analysis, we assumed constant *σ* = 20 nm. However, *σ* ^2^ may change depending on the underlying diffusion dynamics: if molecules diffuse rapidly during the exposure time, the resulting image is blurred, and *σ* becomes larger than the localization error of a stationary molecule, denoted *σ*_0_. This relationship is shown in Eq. (9), which explains the difference between *σ* and *σ*_0_.

Here, we use the exact expression for *σ* to revisit the colormaps shown in Fig. 2 and Fig. 3. From imaging individual mEos3.2 molecules with *t*_*E*_ =20 ms, we found the average photon count to be 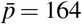. PSF of our microscope setup was measured from imaging surface-immobilized Cy3 molecules (in the same emission channel as photo-converted mEos3.2) and found to be *s*_0_ = 0.256 *µ*m. Based on these values, the localization precision for stationary mEos3.2 would be 20 nm, calculated from 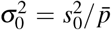. We note that this formula neglects pixelation-related errors from the camera, which are accounted for in more detailed models.^27^ Next, we assumed that 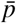 increases linearly with light exposure time, *t*_*E*_.

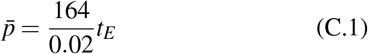

Plugging this expression for 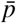 in Eq. (9) as below:

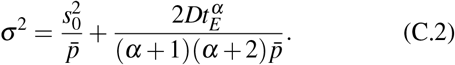

Using this equation, we updated *σ* for each set of *D, α*, and *t*_*E*_ and recalculated the relative error between log(MSD^*′*^) and log(MSD), as shown in Fig. 7. The updated colormaps are largely consistent with those presented in Fig. 2 and Fig. 3, but they additionally capture the negative impact of large *t*_*E*_ on imaging blurring, reflected in increased *σ*, as shown in Fig. 7(b).

**FIG. 7.**
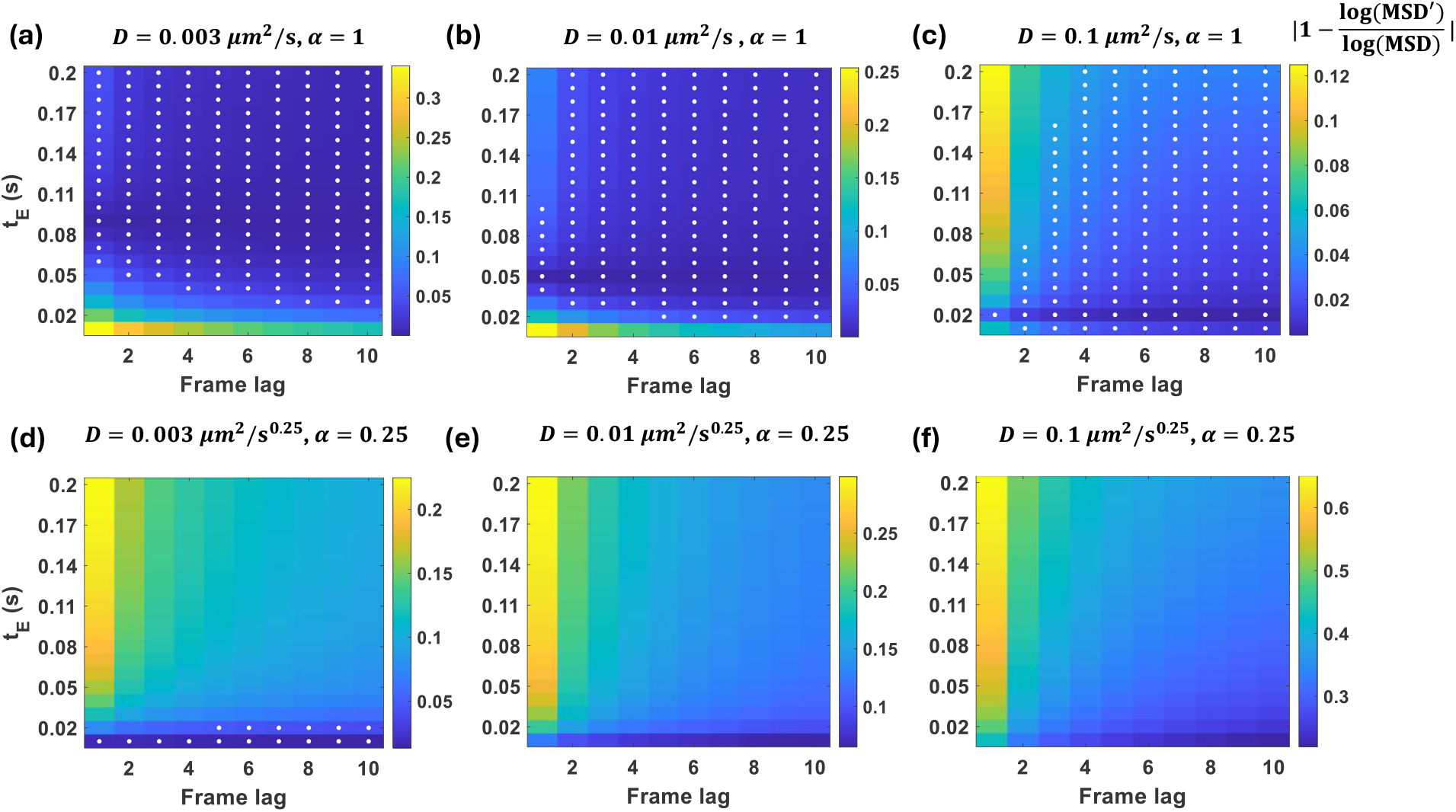
Relative error in the observed MSD (MSD*′*) due to microscopy-induced errors. The observed MSD was computed using Eq. (8) and Eq. (C.2), which considers *σ* as a function of *D, α*, and *t*_*E*_. Diffusion parameter values are indicated in the title of each panel. White dots mark parameter combinations for which the relative error is 5% or less.

